# SARS-CoV-2 and its variants, but not Omicron, induces thymic atrophy and impaired T cell development

**DOI:** 10.1101/2022.04.07.487556

**Authors:** Zaigham Abbas Rizvi, Srikanth Sadhu, Jyotsna Dandotiya, Akshay Binyka, Puja Sharma, Virendra Singh, Vinayaka Das, Ritika Khatri, Rajesh Kumar, Sweety Samal, Manjula Kalia, Amit Awasthi

## Abstract

Pathogenic infections cause thymic atrophy, perturb thymic-T cell development and alter immunological response. Previous studies reported dysregulated T cell function and lymphopenia in coronavirus disease-19 (COVID-19) patients. However, immune-pathological changes, in the thymus, post severe acute respiratory syndrome coronavirus-2 (SARS-CoV-2) infection have not been elucidated. Here, we report SARS-CoV-2 infects thymocytes, depletes CD4+CD8+ (double positive; DP) T cell population associated with an increased apoptosis of thymocytes, which leads to severe thymic atrophy in K18-hACE2-Tg mice. CD44+CD25-T cells were found to be enriched in infected thymus, indicating an early arrest in the T cell developmental pathway. Further, Interferon gamma (IFN-γ) was crucial for thymic atrophy, as anti-IFN-γ antibody neutralization rescued the loss of thymic involution. Therapeutic use of remdesivir (prototype anti-viral drug) was also able to rescue thymic atrophy. While Omicron variant of SARS-CoV2 caused marginal thymic atrophy, delta variant of SARS-CoV-2 exhibited most profound thymic atrophy characterized by severely depleted DP T cells. Recently characterized broadly SARS-CoV-2 neutralizing monoclonal antibody P4A2 was able to rescue thymic atrophy and restore thymic developmental pathway of T cells. Together, we provide the first report of SARS-CoV-2 associated thymic atrophy resulting from impaired T cell developmental pathway and also explains dysregulated T cell function in COVID-19.

---

SARS-CoV-2 which was first reported in Wuhan, China in Dec, 2019 has so far infected around 5.6% of the total world population with around 1.3% mortality rate as on 29^st^ Feb, 2022 (https://covid19.who.int/). While majority of clinical cases report pulmonary pathologies associated with pneumonia and acute respiratory distress syndrome (ARDS), growing number of evidences suggest cardiovascular, gastrointestinal, renal, neurological, endocrinological manifestations of SARS-CoV-2 infection (Chung et al., 2021; Guan et al., 2020; Gupta et al., 2020; Yang and Tu, 2020). Clinical cases of COVID-19 are also characterized by lymphopenia which is defined by T cell lymphodepletion (Tan et al., 2020). In line with this, our clinical data showed a significant depletion of peripheral CD3+ and CD8+ T cells of COVID-19 patients as compared to the healthy control while a decreasing trend was observed for CD4+ T cells (**Fig 1A)**. Changes in the thymus post-acute pathogenic infection have been previously shown to cause lymphopenia as thymus is the site of T cell development and maturation (Liu et al., 2014; Luo et al., 2021). Therefore, we speculated that the changes in the thymocytes developmental pathway could be a driving factor for immune dysregulation as seen in COVID-19. For this purpose, we used acute model of SARS-CoV-2 infection i.e. hACE2-transgenic (Tg) mice expressing humanized ACE2 receptor driven by epithelial cell cytokeratin-18 (K18) promoter. hACE2-Tg mice has been shown to develop acute SARS-CoV-2 infection following intranasal challenge with the live virus associated with rapid loss in body mass leading to mortality by day 6-8 post infection (McCray et al., 2007; Winkler et al., 2020). Consistent with the published reports, we found that hACE2-Tg mice infected with SARS-CoV-2 (B) develops acute COVID-19 pathology characterized by sharp decline in body mass (**Fig S1A & S1B)**, 80% mortality by day 7 and profound presence of lung inflammation and injury as observed by assessing excised lung and hematoxylin and eosin (H&E) stained lung sections by trained pathologist (**Fig S1C-S1G)**. In line with this, there was profound localization of N protein in the infected lungs sections as determined by immunohistochemistry (IHC) corresponding to high N gene copy number (by qPCR) (**Fig. S1H & S1I)**. Characteristic of the active infection, mRNA expression of anti-viral genes which are activated post sensing of viral RNA by toll-like receptors (TLRs) or rig-like receptor (RLRs) and activation of stimulator of IFN genes (STING) such as 2’-5’- oligoadenylate synthetase (OAS)-2 and OAS-3, latent RNase (RNaseL), IFN-induced transmembrane (IFITM) protein, adenosine deaminase acting of RNA-1 (ADAR-1) (Schneider et al., 2014) were all significantly increased in infected lung (**Fig. S1J)**. Interestingly, the spleen and lymph node size of the infected mice at 6 dpi (peak of acute infection) showed profound involution with decrease in mass and live cell population (**Fig S1K-S1O**).

**Figure 1.**
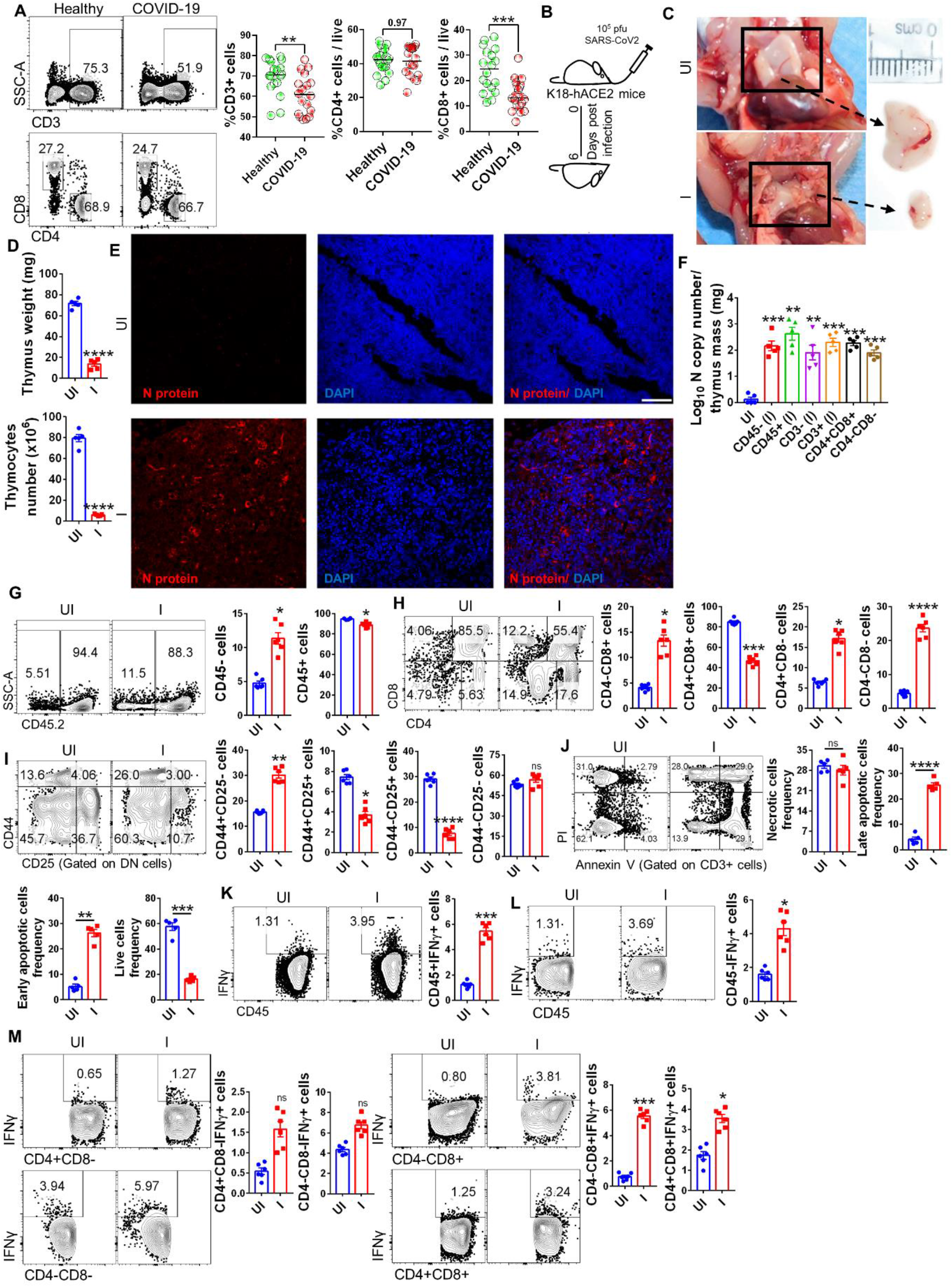
Immuno-pathology of thymus of SARS-CoV-2 infected animalsnn. (A) Immunophenotyping of COVID-19 patients compared to healthy controls (n=18 each). Each dot SSC-A vs CD3, CD4 vs CD8 represents the %frequency of respective cell types per total live cells in PBMCs of an individual subject. Bars represent median values for % CD3 cells/live % CD4 cells/live % CD8 cells/live. **(I-M)** hACE2-Tg mice intranasally (I) infected with 10^5^ pfu SARS-CoV-2 were euthanized at the time points indicated and immuno-pathological changes in the thymus was studied as compared to the uninfected (UI) control. (B) Schematic representation of hACE2-Tg transgenic mice study. The animals were sacrificed on day 6 or on the day they become moribund or otherwise indicated. (C) Representative image of excised thymus along with thymus (D) weight and number. (E) Immunofluorescence microscopy image of thymus section showing SARS-CoV-2 specific N protein (red color), DAPI stain (blue color) and overlay image (DAPI+N protein) acquired at 60X magnification. (F) bar graph showing mean ± SEM of N gene copy for sorted populations of thymus. Representative FACS dot plot and its corresponding bar graph showing percentage frequency mean ± SEM showing (G) SSC-A vs CD45.2 (H) CD4 vs CD8 gated on CD45.2+ cells (I) CD44 vs CD25 gated on DN cells (J) PI vs Annexin V gated on CD3+ cells (K) IFN-γ vs CD45+ (L) IFN-γ vs CD45-and (M) IFN-γ vs different CD4/CD8 sub-populations in the thymus. (D-F) n=5; (B-C, G-M) n=6. Non-parametric t-test using Mann-Whitney test. *P < 0.05, **P < 0.01, ***P < 0.001, ****P < 0.0001.

Emerging studies now show that lymphopenia is strongly correlated to morbidity and mortality and is associated with more than 50% of the adults and 10% of children infected with SARS-CoV-2 (Adamo et al., 2021; Mathew et al., 2020). Lymphopenia is characterized by a substantial decrease in lymphocyte count and reduction in the frequency of peripheral CD4+ T helper and CD8+ T cytotoxic cells (Mathew et al., 2020). In line with this, we observed significantly reduced frequency as well as cell count of CD45.2+, CD3+, CD4+ and CD8+ lymphocytes in the splenocytes of the infected hACE2-Tg mice at 6 dpi as compared to uninfected mice (UI) (**Fig. S1P-S1Q**). Curiously, there was a 3-4 folds upregulation in the frequency and count of DP cells in the splenocytes, which has been shown to be related to escape of DP cells from the thymus (**Fig. S1P-S1Q**). Lymphopenia seen in acute SARS-CoV-2 infected hACE2-Tg mice strongly pointed to dysregulated T cell development in the thymus. In line with this, the thymus of the infected hACE2-Tg mice at 6 dpi showed profound thymic involution (**Fig 1B & C)** with a 7-8 folds decrease in mass and number of live cells (**Fig. 1D**). Next in order to understand the influence of SARS-CoV-2 infection on the thymic developmental pathway we looked at the virus entry and localization into the thymus. Our immunofluorescence data for SARS-CoV-2 specific anti-N protein localization indicated prominent presence of N protein in the thymocytes of the infected hACE2-Tg (**Fig 1E)**. This was a surprising finding since no study has so far reported the expression of humanized ACE2 receptor in the thymus of hACE2-Tg mice. Since cellular injury of pulmonary and extra-pulmonary organs is characteristic pathological manifestation of COVID-19 and is often ascribed to the cellular entry and presence of virus, we evaluated the viral load in different compartments of thymocytes i.e. CD45+, CD45-, CD3+, CD3-, DP, DN cells in the sorted population (**Fig. S2A)**. Our data shows presence of 2-2.5 log10 N gene copy number/ mg mass in all the compartments of thymocytes, suggesting that virus was able to internalize both in CD45-cells (which comprises of thymic epithelial cells) and CD45+ T cells with equal efficiency (**Fig. 1F**).

Next in order to understand whether virus induced thymic involution could result in dysregulation of T cell development we carried out a detailed immunophenotyping for different developmental stages of T cells in the thymus. We found a substantial increase in the frequency of CD45.2-cells, which was accompanied by expansion of CD45.2+ frequency in the infected thymus, however, there was a significant decrease in the cell count of both CD45.2+ and CD45.2-cells, suggesting that the depletion of CD45.2+ cells were higher than that of CD45.2-cells though there was loss of both CD45.2+ and CD45.2-population due to viral infection (**Fig 1G & S2B)**. In line with this, we found 2 folds decrease in CD4+CD8+ double positive (DP) and ∼ 5 folds increase in CD4-CD8-double negative (DN) population with ∼ 2-3 folds increase in single positive (SP) CD4+ or CD8+ cells indicating a profound dysregulation of T cell developmental pathway in infected thymus (**Fig 1H**). Since dysregulated ratio of DP/DN was observed in thymus of infected mice, we made an attempt to understand at which developmental stage of triple negative (TN) population int the thymus is getting affected. Thymic development of TN occurs in 4 distinct stages viz CD44+CD25-(DN1), CD44+CD25+ (DN2), CD44-CD25+ (DN3) and CD44-CD25-(DN4) (Haynes et al., 1999; Liu et al., 2014). We found that thymus of the infected mice showed accumulated frequency for DN1 stage while a decrease in DN2 and DN3 percent frequency was observed for infected samples, suggesting an arrest at the early stage of T cell developmental pathway (**Fig 1I**). We found ∼ 4-6 folds increase in early and late apoptotic cells CD3+ thymocytes in infected as compared to the uninfected control (**Fig 1J**). Similar trends in the induction of early and late apoptosis was found on CD45+ thymocytes, however CD45-thymocytes showed lesser frequency of apoptotic cells suggesting that CD3+ thymocytes are the major depleted population during SARS-CoV-2 infection in mice (**Fig S2C & S2D)**. When time kinetics of infection pathology was studied to understand the early or late phase induction of thymic atrophy, we found that the degree of thymic atrophy was directly proportional to the severity of coronavirus disease-19 (COVID-19) with profound thymic atrophy at day 6 (but not day 3: I_3_ post challenge) (**Fig S2E-S2J)**.

Several mechanisms have been shown to influence thymic atrophy for pathogenic infection such as NK cells activation, increased levels of glucocorticoids as well as heightened IFN-γ secretion by the thymocytes. Previously studies have shown that pathogenic infections causes thymic atrophy associated with apoptosis of thymocytes which could be mediated by IFN-γ induced tissue injury (Barreira-Silva et al., 2021; Liu et al., 2014; Savino, 2006). Our data shows that SARS-CoV-2 infection results in profound elevation (∼5 folds) of IFN-γ by CD45+ cells and to lesser extend (∼2 fold) by CD45-cells (**Fig 1K & L**). Moreover, SP CD8+ cells were found to be the major contributors of IFN-γ production in the thymus with ∼ 6 fold upregulation during infection (**Fig 1M)** which could be one of the driving factors for thymic atrophy as has been earlier reported (Liu et al., 2014). Other pro-inflammatory cytokines such as Granzyme B (GzB), IL-4 and IL-17A was not found to be significantly altered, however interestingly, Perforin-1 (Prf-1) was found to be significantly up-regulated upon infection **(Fig. S2K-S2N)**. Together, we show that SARS-CoV-2 infection in hACE2-Tg mice results in thymic atrophy. Thymic atrophy was characterized by loss of DP population probably due to apoptosis and heightened IFN-γ which could be due to the persistence of virus in the thymus. IFN-γ, produced by both innate and adaptive arms of the immune system, has been earlier shown to be crucial mediator of inflammation and tissue injury during viral infections.

Moreover, IFN-γ is critical in the induction of activation induced cell death (AICD) in T cells (Refaeli et al., 2002). Elevated levels of IFN-γ was one of the key component of cytokine release syndrome in human and animal model (Moore and June, 2020). In fact, our data indicate that a heightened IFN-γ levels in SARS-CoV2-infected hamster and K18-ACE2-Tg mice. In thymus, IFN-γ has been implicated in apoptosis of thymocytes resulting in tissue injury and thymic atrophy (Elfaki et al., 2021; Liu et al., 2014). One previous study have shown that thymic atrophy caused due to influenza A virus could be alleviated by neutralizing IFN-γ (Liu et al., 2021). Since many aspects of pathogenesis and immunological response of influenza A virus and SARS-CoV-2 bear similarity, we speculated that IFN-γ could be one the key mediators of thymic atrophy for SARS-CoV-2 (Flerlage et al., 2021), and thus neutralize IFN-γ functions by using anti-IFN-γ neutralizing antibody. Anti-mouse IFN-γ neutralization was done one day prior and one day post SARS-CoV-2 infection in hACE2-Tg mice in order to neutralize the effector functions of IFN-γ induced upon SARS-CoV-2 infection **(Fig. 2A)**. There was marginal protection in the body weight loss (**Fig S3A)** but no significant decrease in lung viral load in IFN-γ neutralized animals (**Fig S3B)**. However, thymus of the animals receiving anti-IFN-γ antibody neutralization (I+IFN-γ) showed little or no signs of thymic involution in terms of size, mass and number when compared to the thymus from uninfected mice (**Fig 2B-D**). This corroborated with the decreased level of IFN-γ in animals receiving neutralizing antibody (**Fig. 2E-F**). In line with the rescue of thymus size, the percentage frequency of CD45.2+/- cells were restored to normal levels as seen in uninfected control thymus (**Fig 2G**). Moreover, the percentage frequency of DP and DN cells along with the percentage of DN1-DN4 population was also found to recover to their corresponding uninfected percentage frequency (**Fig 2H-I**). Finally, we showed that neutralization of IFN-γ effectively blunted the apoptosis of thymocytes, thus validating that the important and sufficient mediator of thymic atrophy in SARS-CoV-2 hACE2-Tg mice is IFN-γ (**Fig. 2J)**. Since, SARS-CoV-2 in hACE2-Tg mice resulted in profound IFN-γ response, we asked whether IFN-γ induced tissue injury could also occur in C57BL/6 (WT) mice infected with SARS-CoV-2 without necessitating hACE2 receptor dependent virus entry. Remarkably, SARS-CoV-2 infected WT mice did not show any signs of pulmonary pathology or thymic involution as seen in hACE2 mice (**Fig S3C-S3L)**. Together, our findings show that elevated IFN-γ secretion in SARS-CoV-2 infected hACE2-Tg mice leads to tissue inflammation and injury, thereby causing IFN-γ dependent thymic atrophy.

**Figure 2.**
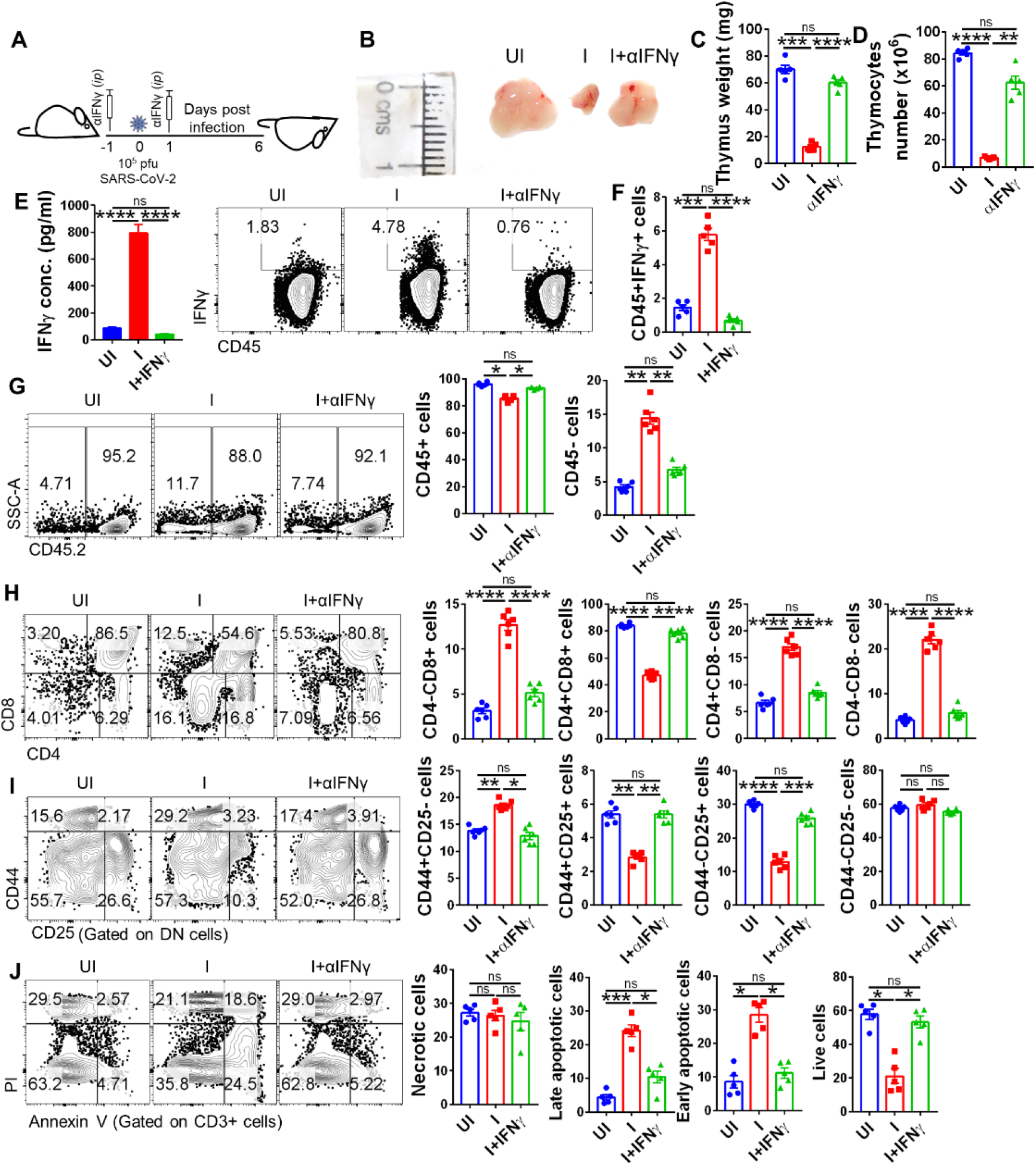
Effect of IFN-γ neutralization on thymic atrophy in SARS-CoV-2 infected hACE2-Tg mice. Neutralizing monoclonal antibody against IFN-γ was used to understand the response of IFN-γ neutralization on thymic atrophy in infected mice. (A) schematic representation indicating schedule for anti-IFN-γ antibody injection. (B) Representative images of excised thymus. (C & D) Bar graph showing mean ± standard error mean (SEM) of thymus wight (mg) and live thymocytes. (E) BALF IFN-γ quantitation by ELISA. Representative FACS dot plot and its corresponding bar graph showing percentage frequency mean ± SEM showing (F) IFN-γ vs CD45 (G) SSC-A vs CD45.2 (H) CD4 vs CD8 gated on CD45.2+ cells (I) CD44 vs CD25 gated on DN cells (J) PI vs Annexin V gated on CD3+ cells in the thymus. (C-E) n=5; (A-B, E-I) n=6. One way-Anova using non-parametric Kruskal-Wallis test for multiple comparison. *P < 0.05, **P < 0.01, ***P < 0.001, ****P < 0.0001.

In the beginning we described that SARS-CoV-2 infection through intranasal route results in virus entry and localization in the thymus. Presence of virus or viral factors have been shown to activate TLR3, TLR7 and TLR8 and RLR which leads to induction of anti-viral genes and cytokines characterized by heightened IFN-γ response (Bonifacius et al., 2021; Moore and June, 2020). This would mean that the use of anti-viral drugs which could reduce the viral burden and reduce viral antigens would also effectively reduce the inflammatory IFN-γ production and hence thymic atrophy. To test this, we used remdesivir (RDV, a prototypic anti-viral drug) which has been shown to have significant efficacy against SARS-CoV-2 infection in clinical cases (Beigel et al., 2020; Wang et al., 2020). Challenged animals receiving remdesivir (I+RDV) treatment significantly reduced lung viral loads and rescued mice from SARS-CoV2 induced pathologies (**Fig 3A, S3M-S3N**). In line with this, RDV treatment in SARS-CoV2 infected mice significantly reduced thymic involution with size, mass and number of live thymocytes restored (approx.) to that of the uninfected animals **(Fig 3B-D)**. Corresponding to this protection, there was profound reduction in the viral load throughout the thymus as assessed by immuno-fluorescence microscopy and qPCR (**Fig 3E-3F**). In addition, animals receiving RDV treatment resulted in alleviation of thymic atrophy immune profile and rescued the normal developmental pathway of thymocytes with restored levels of CD45+/-, DP, DN cells (**Fig 3G-H)**. In line with this, the developmental stages of DN population from DN1-DN4 was restored back to their corresponding uninfected profiles **(Fig. 3I**). Next, we also evaluated the levels of IFN-γ and thymocytes apoptosis as it was important to understand whether RDV mediated rescue of thymic atrophy was operational through clearance of viral load from the thymus or it also resulted in blunted the inflammatory response and related thymic injury. IFN-γ response of CD45+ cells were reduced following RDV treatment and it also resulted in reduced percent frequency of thymocytes entering apoptosis which was expected as viral load and inflammation were earlier shown to be the driving factors for apoptosis of thymocytes **(Fig. 3J-L)**. Together, we establish that reminiscent of IFN-γ, use of RDV (anti-viral drug) could effectively reduce the viral burden could rescue the thymic atrophy induced by SARS-CoV-2 infection.

**Figure 3.**
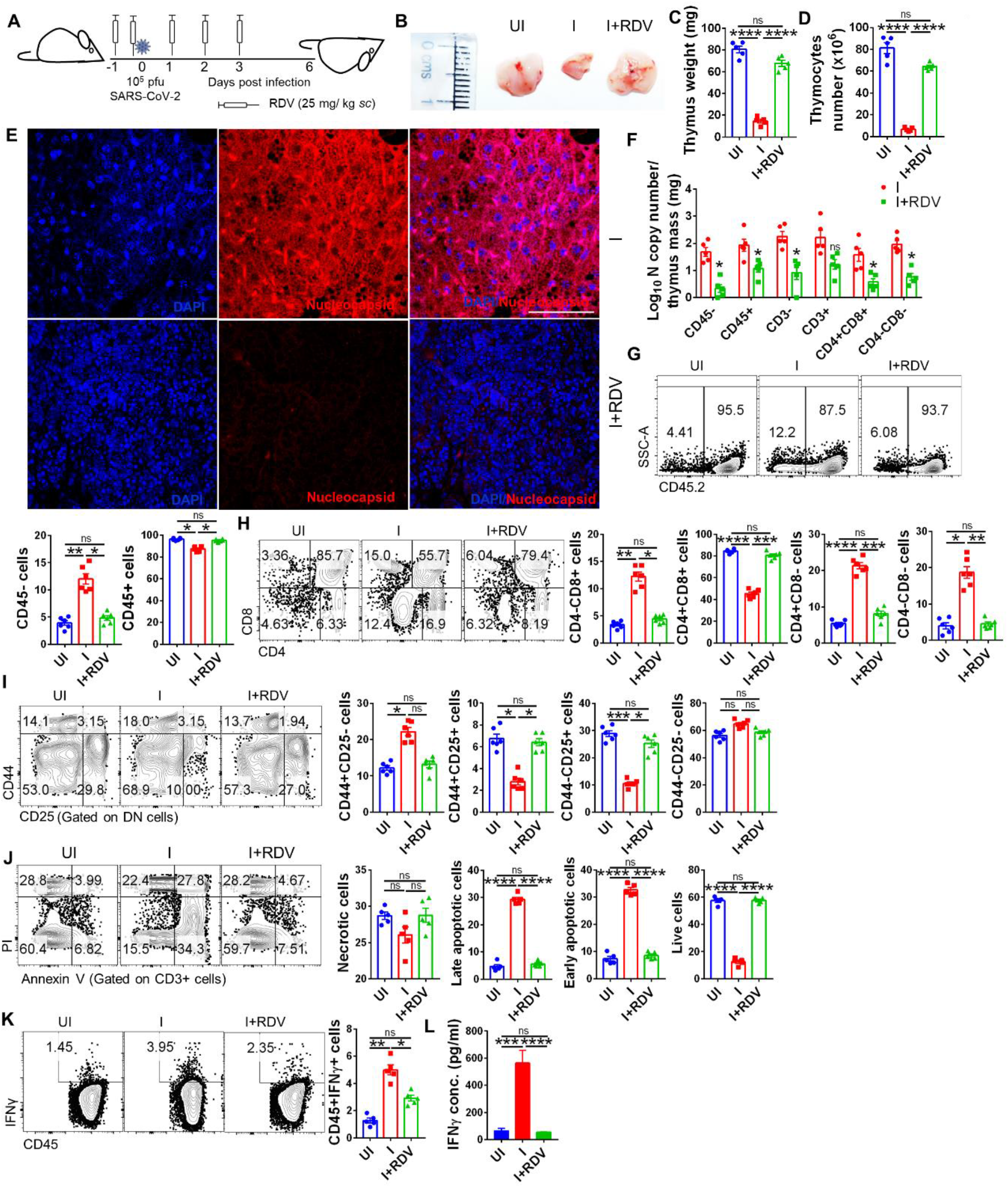
Effect of remdesivir treatment on thymic atrophy in SARS-CoV-2 infected hACE2-Tg mice. Protective efficacy of remdesivir (RDV), used as a prototypic anti-viral drug, against induction of thymic atrophy upon SARS-CoV-2 infection in hACE2-Tg mice. (A) schematic representation showing treatment regime and dose of remdesivir given to SARS-CoV-2 infected mice. (B) Representative images of excised thymus. (C & D) Thymus weight and number shown by bar graph indicating mean ± standard error mean (SEM). (E) Images of immunofluorescence microscopy of thymus section showing SARS-CoV-2 specific N protein (red color), DAPI stain (blue color) and overlay image (DAPI+N protein) acquired at 60X magnification. (F) N gene copy number for sorted thymic populations shown by bar graph mean ± SEM. FACS representative dot plot and its respective bar graph showing percentage frequency mean ± SEM indicating (G) SSC-A vs CD45.2 (H) CD4 vs CD8 gated on CD45.2+ cells (I) CD44 vs CD25 gated on DN cells (J) PI vs Annexin V gated on CD3+ cells (K) IFN-γ vs CD45 in the thymus. (L) Evaluation of IFN-γ from BALF samples by ELISA. (C-F, L) n=5; (A-B, G-K) n=6. One way-Anova using non-parametric Kruskal-Wallis test for multiple comparison. *P < 0.05, **P < 0.01, ***P < 0.001, ****P < 0.0001.

In contrast to hACE2-Tg mice, golden Syrian hamster model mimics mild to moderate coronavirus disease-19 (COVID-19) as observed in majority of clinical cases (Chan et al., 2020). In addition, we had earlier shown that the pulmonary pathology of coronavirus disease-19 (COVID-19) following SARS-CoV-2 infection in hamster peaks at 2-4 days post infection (dpi) with highest lung viral load, and starts to decline by 5-6 dpi with comparatively lower viral load and pulmonary pathology on 7 dpi (Rizvi et al., 2022). Induction of thymic atrophy was evaluated at early and late phase in infected hamsters i.e. 4 and 7 dpi (**Fig S3O**). Though there was no significant difference in the size and mass of the excised thymus at 4 and 7 dpi as compared to that of 0 dpi thymus (**Fig S3P & S3Q**), the number of live thymocytes as measured by trypan blue exclusion dye was significantly lower at 4 dpi (peak of infection) as compared to 0 dpi (uninfected) or 7 dpi (recovery phase of infection), suggesting signs of thymic injury at the peak of infection (**Fig S3R**). Thymus of the 4 dpi hamsters showed 100-500 copy number of viral gene N/ mg of thymus indicating thymic entry of SARS-CoV-2 (**Fig S3S**). We used anti-mouse CD4 (GK 1.5, cross reactive to hamster CD4) and anti-rat CD8 (cross reactive to hamster CD8) both having cross-reactivity with respective CD markers of hamster to study the development of T cells in thymus. Remarkably, we found 12-15% decrease in the DP cells at 4 dpi and 3-5% decrease in DN at 7 dpi as compared to the 0 dpi thymus DP cells. Similar dysregulated ratios were observed for SP CD4 and CD8 cells at 4 dpi. There was a sharp increase in DN cells at 4 dpi with approx. 2-fold increase as compared to 0 dpi thymus suggestive of dysregulated T cell development (**Fig S3T**). In line with this, we found 3-4 folds increase in early and late apoptotic cells at 4 dpi thymus accompanied with approx. 3 folds increase in thymic IFN-γ levels as compared to uninfected control (**Fig S3U & S3V**). Together, our findings show that acute COVID-19, but not moderate COVID-19 leads to severe thymic atrophy.

The original Wuhan strain of SARS-CoV-2 (2019-nCoV) which caused first wave of pandemic in 2020 acquired continuous and considerable mutations in genetic material leading to antigenically different mutations which have been so far characterized into variants of concern (VoC), these VoCs lead to subsequent waves of pandemics across the globe or at different regions of the globe at different time points (Geers et al., 2021). One such major VoC reported for SARS-CoV-2 was beta variant (B.1.351) that was first detected in South Africa with a large number of mutations in the spike region. The two other notable VoC appeared were Delta variant (B.1.617.2) first detected in India in late 2020 and recently reported omicron variant (B.1.1.529) (McCallum et al., 2021, 2022). While both B.1.617.2 and B.1.1.529 have been shown to accumulate large amount of mutations in spike and receptor binding domain (RBD) and evade immune response, but only B.1.617.2 but not B.1.1.529 have been shown to cause severe COVID19 in hACE2-Tg animal model (Halfmann et al., 2022). Since mutations reported in the VoC have been shown to evade immune response we became interested to investigate the effect of VoC challenge B.1.351, B.1.617.2 and B.1.1.529 on the induction of thymic atrophy (**Fig 4A**). Consistent with the previously published reports, mice infected with variants except omicron showed rapid decrease in body mass and with presence of high N-gene copy number (**Fig S4A-S4B**). Our data demonstrate a most profound thymic atrophy in animals challenged with B.1.617.2 variant which closely mimicked the degree of thymic dysregulation of 2019-nCoV both in terms of involution of size, mass and thymocytes number. Interestingly, animals challenged with B.1.1.529 showed marginal thymic atrophy when compared to the original 2019-nCoV strain in line with the lower lung viral load **(Fig. 4B-D)**. Consistent with the thymic atrophy we found significant VoC viral load in all the cellular compartment of thymus which was relatively lower for B.1.1.529 variant (**Fig 4E)**. The profile of developing thymocytes as well as levels of IFN-γ secretion followed the pattern of infectivity with B.1.617.2 showing the highest while B.1.1.529 variant showing lowest dysregulation respectively when compared uninfected control (**Fig 4F-4J & S4C-S4G**). Among the other pro-inflammatory cytokines only Prf-1 was found to be significantly elevated across VoC in CD4/CD8 thymocytes subsets (**Fig S4H-S4K)**.

**Figure 4.**
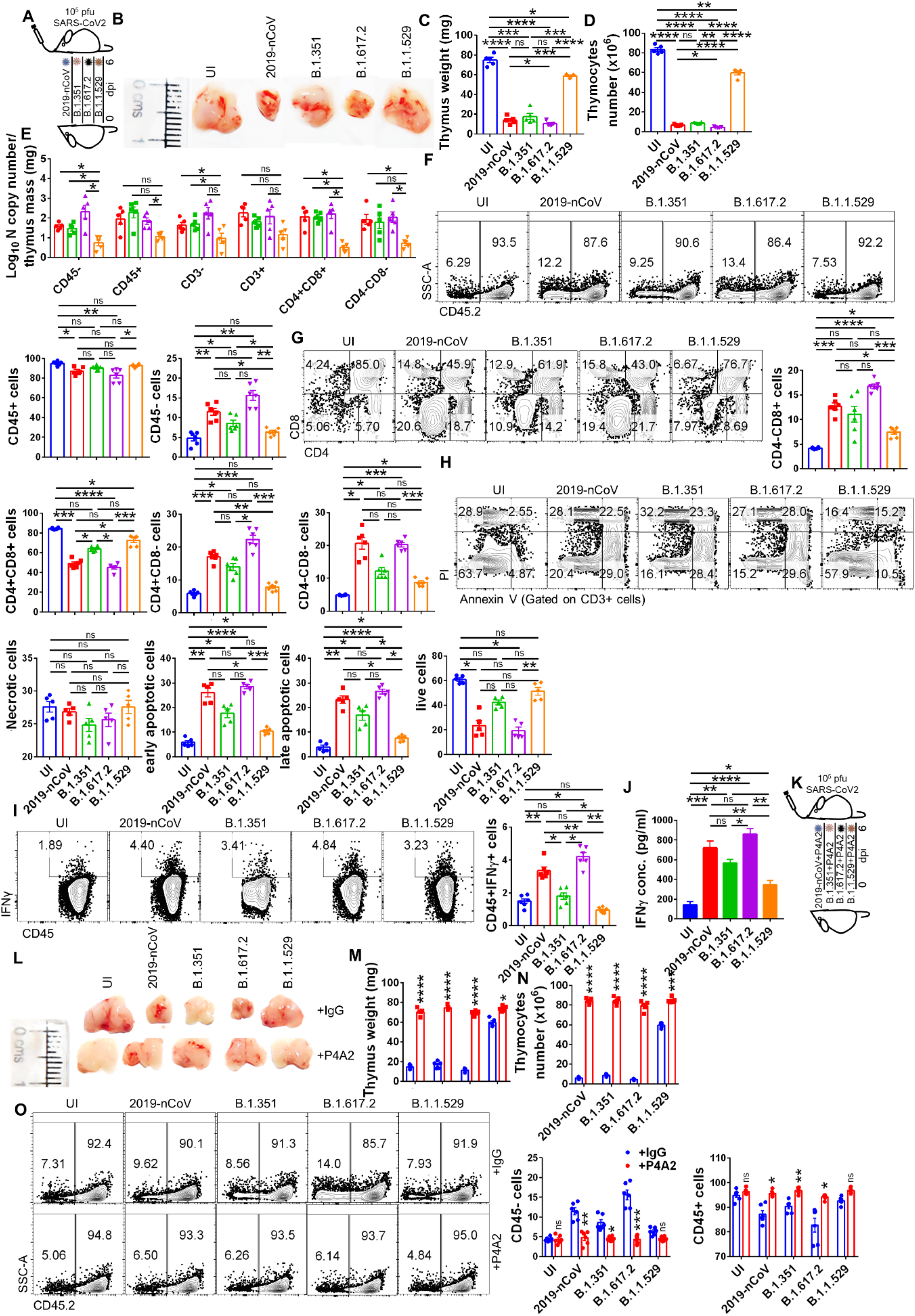
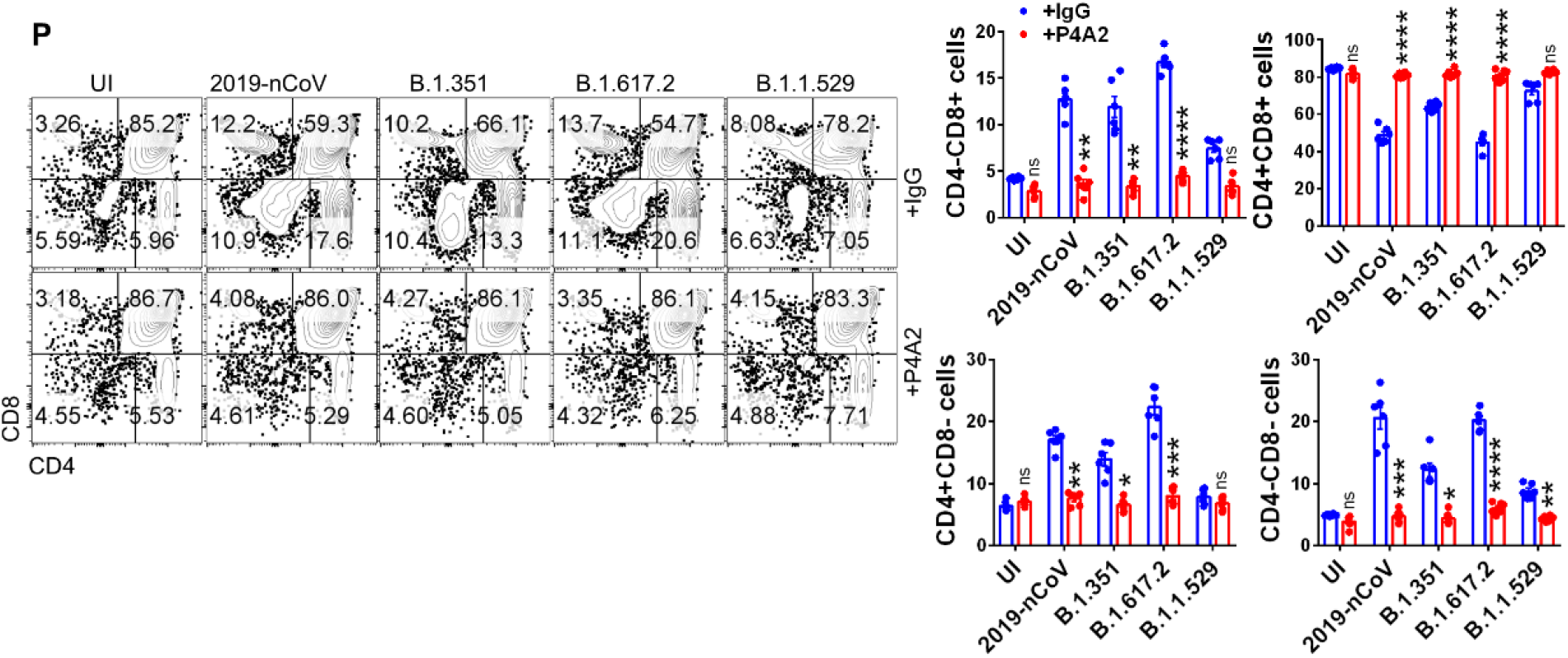
Immuno-pathological changes in thymus infected by SARS-CoV-2 variants and its recovery by broadly SARS-CoV-2 neutralizing monoclonal antibody P4A2. Induction of thymic atrophy in hACE2-Tg mice challenged with SARS-CoV-2 variants such as beta variant (B.1.315), delta variant (B.1.617.2) and omicron variant (B.1.1.529) was studied by intranasal infection 10^5^ pfu/ mice followed by euthanization 6 days post infection or on the day when animals became moribund. (A) schematic diagram for VoC challenged hACE2-Tg mice. (B) Representative images of excised thymus. (C & D) Mean bar graph ± SEM showing thymus wight (mg) and live thymocytes. (E) N gene copy number for sorted thymic populations shown by bar graph mean ± SEM. FACS representative dot plot and its respective bar graph showing percentage frequency mean ± SEM indicating (F) SSC-A vs CD45.2 (G) CD4 vs CD8 gated on CD45.2+ cells (H) PI vs Annexin V gated on CD3+ cells (I) IFN-γ vs CD45 for UI and VoCs in the thymus. (J-O) Therapeutic efficacy of P4A2 broadly SARS-CoV-2 neutralizing antibody in protecting against thymic atrophy by SARS-CoV-2 variants. (J) IFN-γ ELISA from BALF samples. (K) Scheme for VoC challenge in presence or absence of P4A3. (L) Representative image of excised thymus along with (M) thymus weight and (N) number. Representative FACS dot plot and its corresponding bar graph showing percentage frequency mean ± SEM for VoC thymus in presence or absence of P4A2 (O) SSC-A vs CD45.2 (P) CD4 vs CD8 gated on CD45.2+ cells in the thymus. (C-E, J, L-N) n=5; (A-B, F-I, K-L, O-P) n=6. (C-I) One way-Anova (L-O) Two-way Anova using non-parametric Kruskal-Wallis test for multiple comparison. *P < 0.05, **P < 0.01, ***P < 0.001, ****P < 0.0001.

We previously demonstrated that the use of anti-viral drug RDV as an effective prophylactic treatment for rescuing thymic atrophy. We further investigated the therapeutic efficacy of P4A2, a murine monoclonal antibody broadly potent against VoC including Omicron by binding to the receptor binding motif (RBM) of SARS-CoV-2 RBD protein (Kumar et al., 2022). Monoclonal antibodies have previously shown to be effective in neutralizing SARS-CoV-2 in clinical settings and have the advantage of being administered as therapeutic dose post challenge even to immuno-compromised individuals (Brouwer et al., 2020). We found effective neutralization of VoC and alleviation of thymic involution in mice receiving P4A2 antibody (**Fig 4K-N)**. Likewise, the profile of the developing thymocytes along with the masking of IFN-γ secretion was also seen with the therapeutic dosing of P4A2 against all the VoC studied (**Fig 4O-4P)**. Together, we show that the therapeutic dose of P4A2 antibody was effective and sufficient to neutralize VoC challenge and restore the thymic atrophy condition induced by SARS-CoV2 challenge.

In summary, we present the first report of thymic atrophy induced by SARS-CoV-2 infection in both moderate and acute COVID19 model. Lymphopenia and immune dysregulation is a common feature of many pathogenic infection including influenza A and is influenced by the changes in the thymus. Moreover, thymus is the primary lymphoid organ and is central to the immune hemostasis. Through our study, we found that lymphopenia associated with SARS-CoV-2 infection could be attributed to thymic dysregulation and thymic atrophy. Since lymphopenia is known to persist long after COVID19 infection in clinical cases, we believe that thymic atrophy could be one of the important contributors of lymphopenia and may contribute to the alteration in peripheral T cell receptor repertoir. These findings may provide a paradigm change in our understanding of how immune-response (especially T cell response) is modulated during COVID-19 and may provide novel mechanism for designing vaccine candidates since thymic atrophy had been shown to result in loss of TCR repertoire.

## Acknowledgments

Financial support was provided to the AA laboratory from THSTI core, Translational Research Program (TRP), BIRAC grants (BT/CS0054/21 and BT/CTH/0004/21) Department of Biotechnology (DBT) and DST-SERB. ZAR is supported by intramural funding (THSTI). Immunology Core and FACS facility for providing support in experimentation. We acknowledge SAF and infectious disease research facility (IDRF) for its support. ILBS bio-bank for support in histological analysis and assessment. RCB microscopy facility for microscopic examination of the histology slide. We acknowledge the technical support of Manas and Sandeep. SARS-CoV-2 and its variants were deposited by the Centres for Disease Control and Prevention and obtained through BEI Resources, NIAID, NIH: SARS Related Coronavirus 2, Isolate USA-WA1/2020, NR-52281.

## Author Contributions

Conceived, designed and supervised the study: AA; Designed and performed the experiments: ZAR; ABSL3 experiment: ZAR, SS, JD; FACS: ZAR, SS, JD; qPCR: ZAR; ELISA: ZAR; Floursecence microscopy: PS, MK; Antibody P4A2: RK; Analyzed the data: ZAR; Contributed reagents/materials/analysis tools: AA; Wrote the manuscript: ZAR, AA.

## Data availability statement

All data needed to evaluate the conclusions in the paper are present in the paper and/or the Supplementary Materials

